# Metabolic potential, ecology and presence of endohyphal bacteria is reflected in genomic diversity of Mucoromycotina

**DOI:** 10.1101/2020.11.16.384453

**Authors:** Anna Muszewska, Alicja Okrasińska, Kamil Steczkiewicz, Olga Drgas, Małgorzata Orłowska, Urszula Perlińska-Lenart, Tamara Aleksandrzak-Piekarczyk, Katarzyna Szatraj, Urszula Zielenkiewicz, Sebastian Piłsyk, Ewa Malc, Piotr Mieczkowski, Joanna S. Kruszewska, Przemysław Bernat, Julia Pawłowska

## Abstract

We describe the genomes of six Mucoromycotina fungi representing distant saprotrophic lineages within the subphylum (i.e. Umbelopsidales and Mucorales). We selected two *Umbelopsis* isolates from soil (i.e. *U. isabellina, U. vinacea*), two soil-derived *Mucor* isolates (i.e. *M. circinatus, M. plumbeus*), and two Mucorales representatives with extended proteolytic activity (i.e. *Thamnidium elegans* and *Mucor saturninus*). We complement genome analyses with a description of their digestive capabilities, their cell wall carbohydrate composition, and total lipid profiles. Finally, we link the presence of endohyphal bacteria with observed characteristics.

One of the genomes, *Thamnidium elegans*, harbours a complete genome of an associated bacterium classified to *Paenibacillus* sp. This fungus displays multiple altered traits compared to remaining isolates regardless of their evolutionary distance. *T. elegans* has expanded carbon assimilation capabilities particularly efficiently degrades carboxylic acids, has a higher diacylglycerol: triacylglycerol ratio and phospholipid composition suggesting a more rigid cellular membrane.
Comparison of early-diverging Umbelopsidales with evolutionary younger Mucorales points at several differences particularly in their carbon source preferences and encoded carbohydrate repertoire. All tested Mucoromycotina shares features including the ability to produce 18:3 gamma-linoleic acid and fucose as a cell wall component.

**Author Summary:** In our paper, we report on the genomic sequences of six Mucoromycotina strains and an associated bacterium from *Paenibacillus* genus. Mucoromycotina are often studied in pathogenic context albeit their basic biology remains understudied. This manuscript expands on the collection of currently sequenced Mucorales and Umbelopsidales, including the first sequenced *Thamnidium* isolate, which was sequenced together with a *Paenibacillus* bacterium. The interaction with a bacterial partner alters the metabolism, cell membrane composition but not the exoskeleton of the fungus. The associated bacterium provided multiple enzymes that significantly expanded the digestive capabilities of the fungal host. Parallel sequencing and phenotyping of Mucorales and Umbelopsidales enabled us to look at the differences of both lineages within Mucoromycotina. We demonstrate that the predicted digestive capabilities are in line with experimental validation. Based on the cell wall composition data and genomic underpinnings of carbohydrate metabolism we were able to confirm the universal presence of fucose in Mucoromycotina cell walls. Fatty acid, phospholipid and acylglycerol composition support the usage of 18:3 gamma-linoleic acid as a chemotaxonomic marker of Mucoromycotina and corroborate TAG as a dominant storage lipid in these organisms.

Genomic features, digestive capabilities, fatty acid composition differ between Mucorales and Ubelopsidales pointing at subtle but significant changes in the course of Mucoromycotina radiation.

## Introduction

Mucoromycotina subphylum comprises three orders: Umbelopsidales, Endogonales and Mucorales [1]. While Umbelopsidales and Mucorales group mostly saprotrophic fungi living in the soil, on dung and litter, Endogonales are known to establish symbiotic interactions with plants [2]. The ancestors of extant Mucoromycotina were among the first colonizers of land. These early-branching fungi possess all of the traits commonly acknowledged as distinctive characteristics of the fungal kingdom such as apical growth, presence of ergosterol in the membranes and a cell wall made of chitin and beta-glucan [3]. Yet, their cell wall differs from ascomycetes and basidiomycetes by the presence of fucose and high amounts of N-acetylglucosamine and glucuronic acid [4,5] which is typical for other Opisthokonta rather than fungi. Mucoromycotina representatives form a fast-growing mycelium of haploid hyphae. The sexual phase includes the fusion of two gametangia and formation of a resting spore called zygospore [1].

Apart from decomposing organic matter as saprotrophs, some Mucoromycotina are capable of forming mutualistic and mycorrhiza-like associations with Haplomitriopsida liverworts [6,7]. Others are parasites of plants and animals [8,9]. The ecology of Mucoromycotina is poorly studied, which hinders understanding of their role in the ecosystem [10].

The best-studied order of Mucoromycotina is Mucorales. Its representatives are perceived mostly as a source of food spoilage, rotting crops and decomposing litter [11]. Due to their efficient plant matter degrading potential, Mucorales are used in various food production processes like the fermentation of soya beans known in the traditional Asian and African cuisine as tempeh [12] or cheese making [13]. Others are employed as bio producers of polyunsaturated fatty acids (eg. *M. circinelloides*), β-carotene (eg. *Blakeslea trispora, M. circinelloides*, and *Phycomyces blakesleeanus*) and diverse hydrolases [14–16]. Certain Mucorales (e.g. *M. indicus*) are also able to produce ethanol [17] while others (e.g. *Rhizomucor miehei*) are commercially used for lipase production [18][11]. Mucorales lipid composition includes gamma-linolenic fatty acid 18:3 [19] which is found mostly in plants [20], algae [21] and not in Dikarya which produce the n-3 isomer of the C18 trienoic fatty acid alpha-linolenic acid.

Some Mucorales are involved in life-threatening, opportunistic infections with mortality rates reaching up to 40-80% [22]. There are general traits which predispose microorganisms to become opportunistic pathogens, e.g. thermotolerance and ability to evade immune cells. Many non-pathogenic fungi are adapted to higher temperatures due to living in decomposing organic matter, often warmed up due to rotting processes [23,24], which allows them to survive inside the animal warm-blooded body.

Recently, endohyphal bacteria have been described in a few different fungal lineages of Mucoromycota (i.e. Mucoromycotina and 2 other closely related subphyla: Glomeromycotina and Mortierellomycotina). *Mycoplasma-related* endobacteria (MRE) were detected in Endogonales [25], Glomeromycotina [26], and Mortierellomycotina [27]. *Burkholderia-related* endobacteria (BRE) were identified in Mortierellomycotina, Mucorales and Gigasporales representatives [28]. *Paraburkholderia*-related endobacteria (PRE) were discovered in Mucorales, Umbelopsidales, and Mortierellomycotina representatives. Lately, information about two other endosymbionts of *Rhizopus microsporus* has been published: *Ralstonia pickettii* as an enabler of fungal invasion during infection [29] and *Stenotrophomonas* sp. whose role has not yet been established [30]. The activity of endohyphal and fungi-associated bacteria involve (but is not limited to) change in lipid metabolism [31–33], change in morphology of the fungus [31,32,34], control of the hosts’ sexual [35], and asexual reproduction [36].

Here we describe the genomes of six Mucoromycotina species representing separated saprotrophic lineages within the subphylum (i.e. Umbelopsidales and Mucorales). Mucoromycotina are often studied in pathogenic context albeit their basic biology remains understudied. We selected two *Umbelopsis* isolates from soil: *U. isabellina* and *U. vinacea*, two soil-derived *Mucor* isolates: *M. circinatus* and *M. plumbeus*, and two Mucorales representatives: *Thamnidium elegans* and *M. saturninus* with proteolytic capabilities enabling them to colonize dung and animal substrate [37]. We complement genome analysis with a phenotypic description of their digestive potential, their cell wall carbohydrate composition, and total lipid profiles. Finally, we link the presence of endohyphal bacteria with observed characteristics.

## Results

### Genome assembly and annotation

Obtained assemblies showed diverse levels of fragmentation depending on genome size and abundance of repeats. Both *Umbelopsis* genomes assembled into fewer than 200 scaffolds whereas *Mucor* assemblies were significantly more fragmented (**Table 1**).

**Table 1.** Summary of genome sequence characteristics.

|  | *Umbelopsis isabellina* | *Umbelopsis vinacea* | *Mucor circinatus* | *Mucor plumbeus* | *Mucor saturninus* | *Thamnidium elegans* | *Paenibacillus* sp. |
| --- | --- | --- | --- | --- | --- | --- | --- |
| genome size [Mb] | 22 | 23.2 | 55.7 | 49.5 | 40.3 | 31 | 6.9 |
| genes | 9030 | 9241 | 14148 | 11690 | 13020 | 10765 | 5954 |
| tRNAs | 129 | 134 | 194 | 274 | 262 | 191 | 119 |
| TE number | 37 | 59 | 1681 | 869 | 1166 | 541 | - |
| % GC | 42 | 43.09 | 31.89 | 31.1 | 36.59 | 33.15 | 44.61 |
| % repeats | 7.85 | 7.81 | 28.24 | 43.3 | 32.52 | 18 | - |
| contigs | 82 | 175 | 16245 | 8313 | 7254 | 3243 | 285 |
| coverage (x) | 60 | 68 | 26 | 28 | 31 | 61 | 8 |
| N50 | 1335078 | 1328092 | 125937 | 49105 | 159134 | 100281 | 129168 |
| L50 | 7 | 7 | 116 | 301 | 68 | 89 | 16 |
| BUSCOS % | 98.3 | 97.6 | 96.9 | 97.2 | 98.7 | 94.5 | 85.4 |

Genome completeness was verified using single copy fungal orthologous genes searched by BUSCO [38]. Only a few BUSCOSs were duplicated from 27 in *U. isabellina* up to 57 in *M. saturninus* suggests that sequenced genomes were not duplicated. The number of predicted genes, repeats and transposable elements (TEs) content correlates positively with genome size.

#### Repetitive elements

The largest group of TEs found in the genomes of Mucorales, with more than 200 copies per genome are class II elements belonging to Tc1/Mariner superfamily (**Supplementary Table S1**) but other DNA-repeats were also ubiquitous. EnSpm, PIF/Harbinger, hAT-Ac MuLE, Merlin, PiggyBac were present from one up to a hundred copies in all sequenced Mucorales. EnSpm, PIF/Harbinger, hAT-Ac and MuLE are ubiquitous in fungi in general, whereas Merlin elements are characteristic for basal fungal lineages and were apparently lost in Dikarya [39]. Unlike Mucorales, Umbelopsidales genomes had just a few copies of Tc1/Mariner, hAT-Ac, and a single Ginger element. Class I retrotransposon landscape is dominated by LTR retrotransposons from Ty3/Gypsy and LINE/L1 elements which were found in all genomes except for *U. vinacea* which lacks the L1s. Surprisingly the omnipresent Ty1/Copia elements were identified only in the genome of *M. plumbeus* (two copies). Also, all sequenced species had LTR/DIRS elements, characteristic for their tyrosinase integrase and are absent from Dikarya [40]. Mucor genomes harbour also numerous Helitrons, known for their rolling circle replication mechanism. The presence of Helitrons has been observed in other *Mucor* species previously [41]. Noteworthy, Helitrons often hijack neighbouring genes and are efficient vectors of HGT in other fungi [42]. Overall, *Umbelopsis* genomes contained ten-fold less transposable elements per genome compared to Mucorales, which is expected taking into account the differences in genome size.

#### RNAi and other defence mechanisms

Fungal genomes are usually protected *via* diverse mechanisms that include fungal repeat-induced point mutation (RIP), methylation induced premeiotically (MIP), meiotic silencing of unpaired DNA (MSUD) and quelling. It has been described that Mucormycotina possesses RNAi pathway components [43] which are also important for fungus biology [44,45]. We scanned the sequenced genomes with HMM profiles of RNAi core enzymes, namely Dicer, Argonaute and RdRP, and demonstrate that all are present in all six fungal genomes, with duplication of RdRP in all Mucorales (from three to five copies per genome) and a single copy in both *Umbelopsis* isolates (**Supplementary Table S2)**. *U. vinacea, M. circinatus, M. plumbeus* harboured duplicated Argonaute proteins.

Self-non-self and the fungal immune system based on Nucleotide Oligomerization Domain (NOD)-like receptors (NLRs) could be identified neither in the six genomes described in this work nor in 14 predicted Mucormycotina proteomes available at NCBI. The typical NLR central domains NACHT or NB-ARC seem to be absent from those genomes.

### Detection of *Paenibacillus* bacteria in *Thamnidium* genome

Initial *Thamnidium* assembly contained genome fragments characterized by two clearly distinct GC content ratios and for that reason was additionally analysed as a metagenome. It is composed of two easily distinguishable fractions, one belonging to a presumed fungal host and the latter to an associated bacterium representing Paennibacillales, Firmicutes. Fungal and bacterial genomes were re-assembled separately. Remaining five Mucoromycotina assemblies did not contain significant amounts of sequences with sequence similarity to non-Mucoromycotina taxa.

#### *Paenibacillus* features

The bacterial genome is moderately complete (BUSCO score 85.7%) and encodes 5815 genes. Its closest relative, *Paenibacillus* sp. 7523-1, despite DDH similarity above 70% threshold for two out of three DDH calculation formulas implemented in TYGS (67.2; 88.3; 72.9), differs in GC content by 2.35% which supports the separation of the newly sequenced strain as a new species (**Supplementary Figure S1, Supplementary Table S11**). A phylogenetic tree inferred from 16S RNA of related isolates shows its proximity to *P. illinoensis* isolates despite differences in GC content (**Fig.1.**). GC content of the identified *Paenibacillus* genome could be altered by misplaced fungal reads. Despite discarding all reads mapping on eukaryote genomes when assembling the *Paenibacillus* genome it is still possible that some fungal fragments, not similar to other eukaryotic sequences, could not be filtered out.

**Figure 1.**
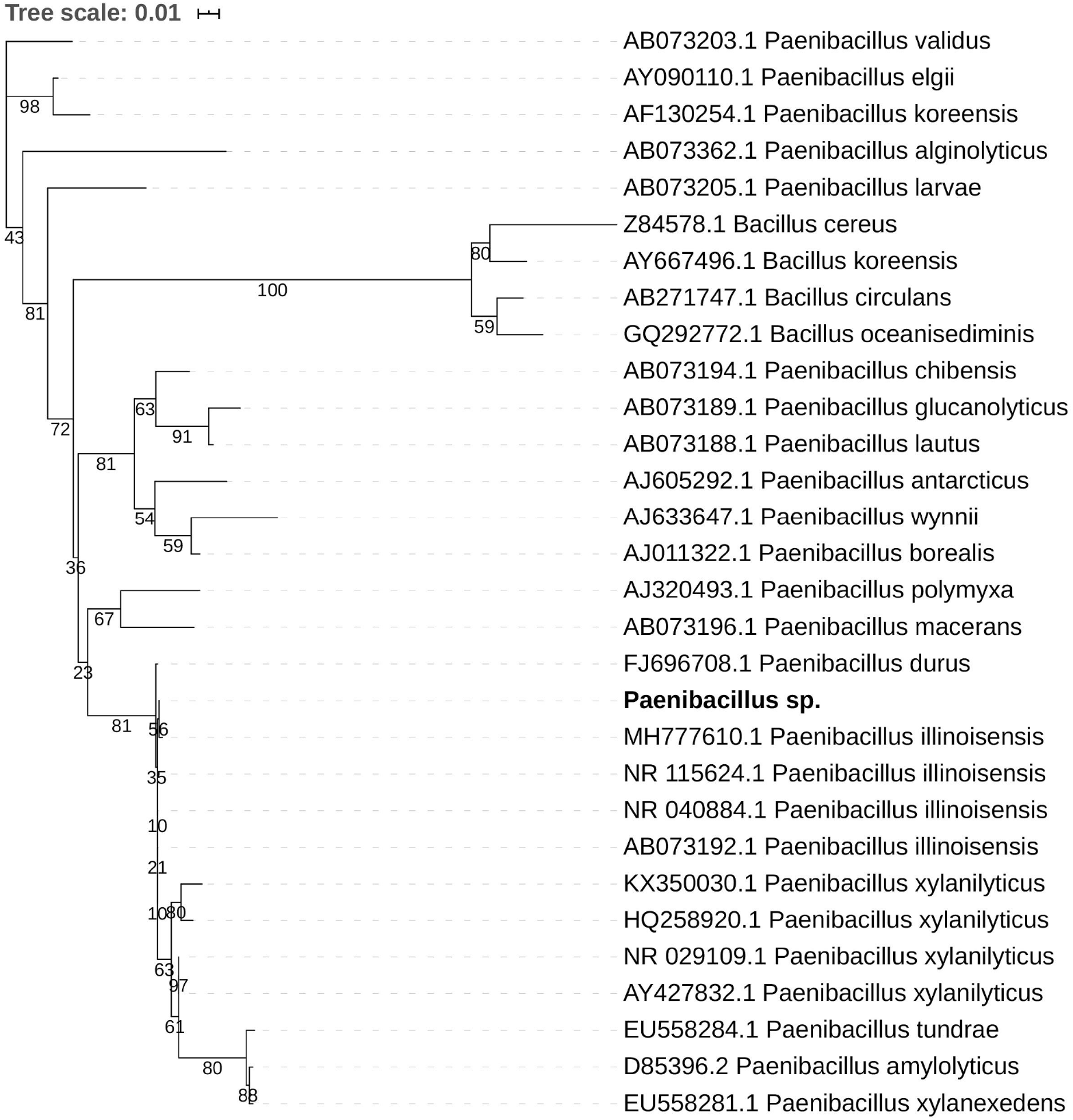
Maximum likelihood phylogenetic tree of 16S rRNA shows the sequenced strain (bold font) groups with *P. illinoisensis* isolates. Branches with support values below 50 were collapsed, the tree was rooted with *Bacillus* sequences.

The bacterial genome has only one complete antimicrobial locus, a rifamycin-inactivating phosphotransferase RphD, identified by scanning using ABRicate. We found a partial beta-lactam resistance pathway lacking the central beta-lactamase (four copies of beta-N-acetylhexosaminidase, BlaI family transcriptional regulator, oligopeptide transport system ATP-binding protein from oppA to oppF, penicillin-binding protein 1A and 2A). Additionally, the *Peanibacillus* genome encodes a single M56 peptidase and 15 proteins bearing an S12 domain. Family M56 includes BlaR1 which is the antirepressor in beta-lactamase operon [46]. Family S12 groups carboxypeptidases B vital for cell wall synthesis and remodelling, as well as betalactamases [47]. Taken together, all these traces point at a potential beta-lactam resistance of *Paniebacillus*. Glycopeptide antibiotic resistance may also be present in some form since partial vancomycin resistance operon was also identified, once more only regulation and accessory components are preserved. The possible activity of these proteins and their relevance for resistance is not known.

The *Paenibacillus* genome harbours vitamin biosynthesis pathways e.g. cobalamin (B12), riboflavin (B2), menaquinone (K2) and thiamine (B1). It also has all the necessary genes for molybdenum cofactor synthesis and manipulation. *Paenibacillus* has many quorum sensing genes and those coding for flagellar and mobility proteins. Its metabolic potential is described below in parallel with its fungal host and remaining fungal genomes.

The *Paenibacillus* genome encodes 3 copies of Peptidase_U32 (PF01136), a collagenase that may facilitate meat degradation by *Thamnidium elegans* which is one of the few fungi known for this property [48]. This protein is found mostly in Firmicutes and is absent from eukaryotic genomes. The U32 collagenases are considered as virulence factors in animal-infecting bacteria [49]. The U32 collagenase (PrtC) together with urease subunit alpha (UreB) are parts of the *Helicobacter pylori* arsenal used in epithelial cell invasion [50]. *Paenibacillus* sp. genome harbours all three urease subunits (alpha, beta and gamma). Moreover, we identified genes coding other urea processing enzymes including: cyanuric acid amidohydrolase, biuret amidohydrolase (BiuH), urea carboxylase and two copies of allophanate hydrolase (AtzF). The *Paenibacillus* genome encodes also one copy of ulilysin Peptidase_M43 (PF05572) with possible gelatinase function [51], and 33 copies of Peptidase_M23 (PF01551), which includes mostly bacterial peptidoglycan hydrolases but also prokaryotic collagenases [52].

### Genes coding for CAZymes, proteases and transport-related proteins

#### Proteases

The overall profile of encoded proteases is very similar in all sequenced fungi (**Fig.2.**), with high numbers of encoded pepsin-like A01A peptidases, FtsH-like M41 peptidases, M48 peptidases active on di- and tripeptides, proteasome peptidases T1, lysosomal peptidases C26, and ubiquitin-specific proteases C19. These proteases, except for pepsin, contribute to intracellular protein turnover and regulation. Pepsin and subtilisin proteases are found in high copy numbers in all of the genomes pointing at a high degradation potential of these saprotrophic organisms. In all analyzed fungal genomes, we found an expansion of the C44 family. Proteins from this family are homologs of glutamine-fructose-6-phosphate transaminase (GFPT) and its precursor. GFTP is known to control the flux of glucose into the hexosamine pathway which plays a crucial role in the regulation of chitin synthesis in fungi [53]. There is a two-fold expansion of S8A serine proteases in Mucorales when compared to Umbelopsidales (**Supplementary Table S3)**. The same observation applies to family I4 which groups inhibitors of S8 peptidases. Umbelopsidales possess several families of cysteine peptidases (C15, C40, C110) and metalloproteases (M14A and M20A) absent from Mucorales genomes. The metalloproteases are likely involved in protein degradation whereas the cysteine peptidases have no obvious function in fungi.

**Figure 2.**
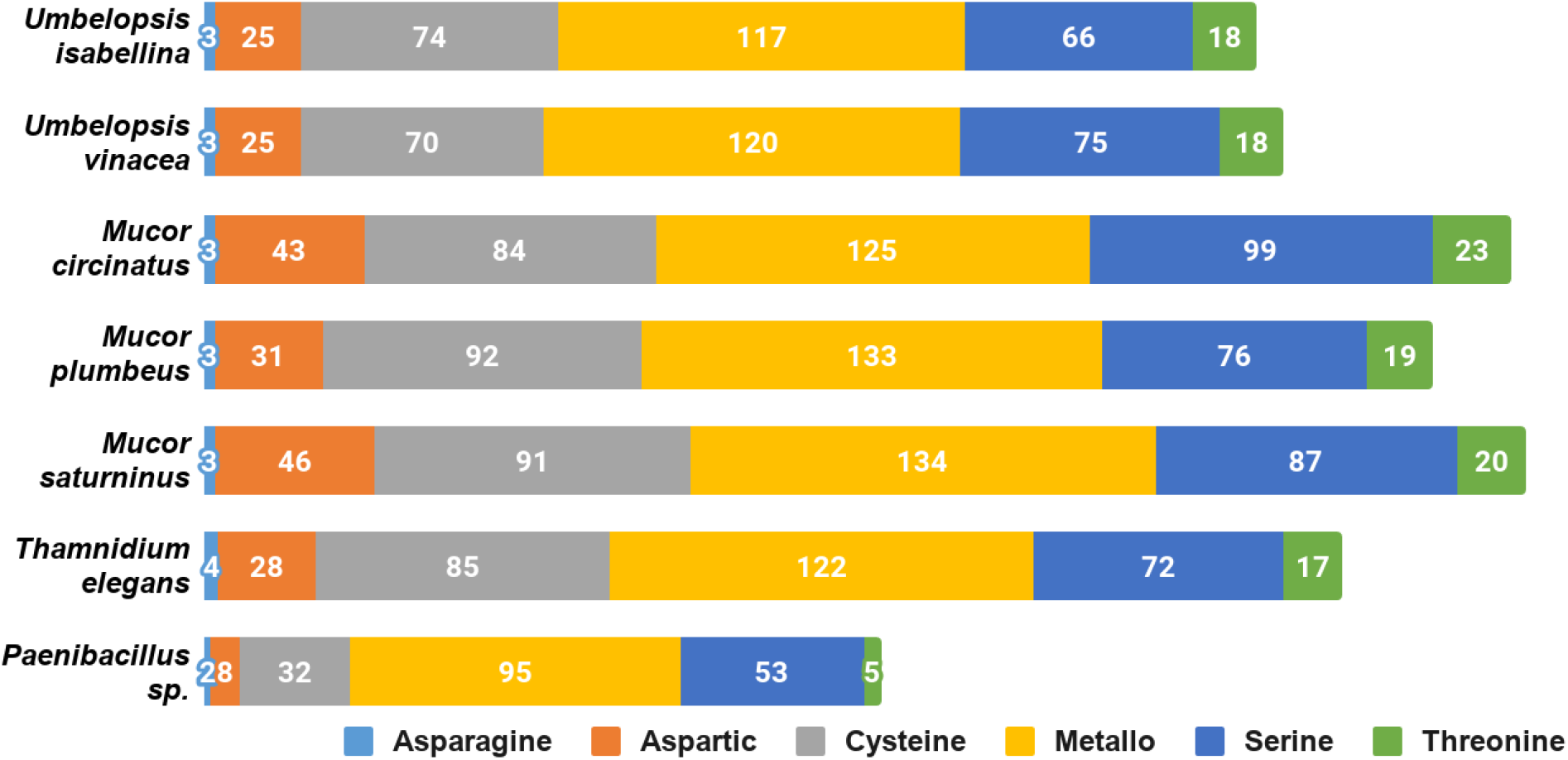
Distribution of genes encoding particular catalytic types of proteases among sequenced isolates.

Among sequenced Mucorales, the average number of protein-coding genes per peptidase family is elevated, which could be a consequence of the whole genome duplication (WGD) described in *Mucor* and *Phycomyces* by Corrochano and coworkers [54]. However, BUSCO results show a low level of duplicated genes in all of the isolates. The genome of *Thamnidium* stands out as it encodes fewer proteases than remaining Mucorales. However, it may benefit from enzymes provided by its endohyphal bacterium which has genes encoding meat crumbling collagenase U32 and plenty of typically bacterial proteases representing families A36, A25, S15, S55, S66, S51, M29, and U57.

#### Carbohydrate active enzymes

All six genomes encode an extensive repertoire of carbohydrate-active enzymes (**Fig.3.**). A high number of glycohydrolases is present exclusively in the genomes of *Umbelopsis* spp. and absent from all four Mucorales; these are GH1, GH2, lysozymes (GH24 and GH25), bacterial β-xylosidase GH52, endo-β-1,4-galactanase GH53, β-1,3-glucanase GH55, GH71, rhamnosidase GH78, α-mannosidase GH92, exo-α-L-1,5-arabinanase GH93, α-L-fucosidase GH95 and peptidoglycan lytic transglycosylase (GH104), enzymes that hydrolyze fructose-containing polysaccharides such as inulinases (GH32), endoglucanase like cellulase and endomannanase (GH5), phosphorylases (GH65), β-glucuronidases (GH79) and glucuronyl hydrolases (GH88) (**Supplementary Table S4)**. All studied fungi have numerous copies of enzymes potentially involved in nutrient degradation and metabolite modification, for instance: α-amylase (GH13), β-amylase (GH14), feruloyl esterase/acetyl xylan esterase (CE1), cholinesterase (CE10). All studied genomes have also an acetyl esterase that liberates acetic acid from xylo-oligomers (CE16). All also have homologs of GH13 α-amylases, but only Umbelopsidales have representatives of the GH13_40 subfamily. This subfamily seems to group exclusively eukaryotic α-amylases. Surprisingly, *Umbelopsis* spp. have also PL12 lyase, a bacterial enzyme used to access metazoan debris by specifically cleaving heparan sulfate-rich regions of acidic polysaccharides which are found in connective tissues [55]. Other typically bacterial enzymes found here are β-xylosidase GH52 and peptidoglycan lytic transglycosylase GH104. This repeated presence of bacterial enzymes may be a result of a horizontal gene transfer event from either endohyphal or free-living bacteria to Umbelopsidales or its ancestor. We also observed an expansion of GH16 hydrolases and hexosaminidase (GH20) in Umbelopsidales compared to Mucorales. On the other hand, *Umbelopsis* spp. lack multiple enzymes belonging to several Cazy families, like acetyl xylan esterase (CE6), cellulose-binding modules (CBM6 and CMB9), chitin-binding module (CBM14), cell wall-related α-1,6-mannanase (GH76) and endo-β-1,4-mannanase (GH134).

**Figure 3.**
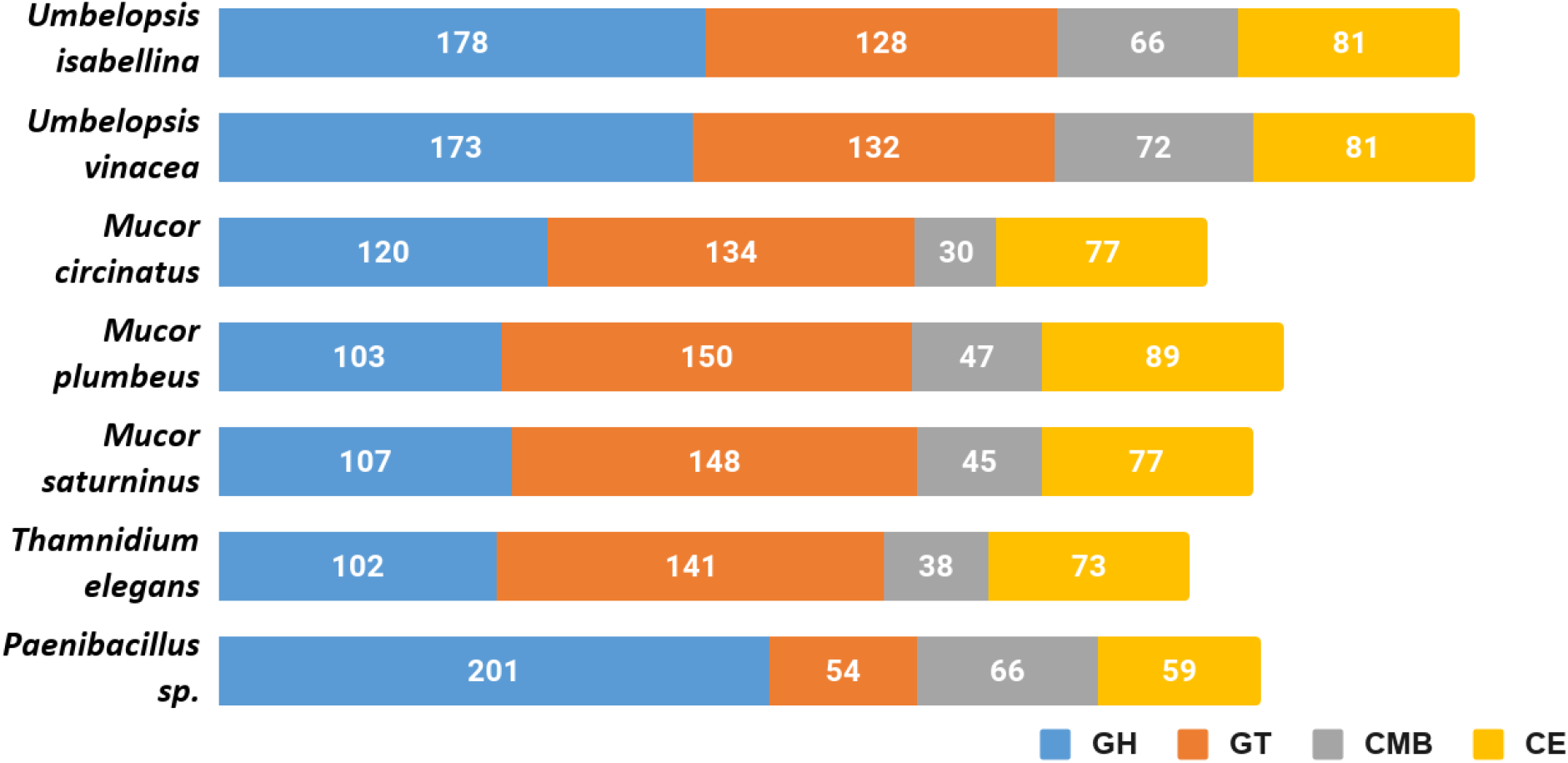
Cazyme distribution in the sequenced genomes. Abbreviations: GH-glycoside hydrolases, GT-glycosyltransferases, CMB-carbohydrate-binding modules, CE-carbohydrate esterase.

Noteworthy, all six fungi have expansions of protein families involved in cell wall remodelling. These include: chitooligosaccharide deacetylase (CE4), chitinase (GH18), chitin synthase (GT2), α-1,2-mannosyltransferase (GT15), β-1,3-N-acetylglucosaminyltransferase / β-1,3-glucuronyltransferase (GT49). Additionally, we found expansions of chitin or peptidoglycan (CBM50) binding modules and additional, broad range binding module (CBM13). Protein glycosylases are also particularly abundant in sequenced taxa and include homologs of Drosophila O-fut2 belonging to protein O-α-fucosyltransferase (GT68) and glycoprotein α-1,3-L-fucosyltransferase (GT10). The latter is encoded by 3 *Mucor* taxa and other known Mucoromycotina but not by *Umbelopsis* spp. Fucosylation may additionally mask Mucorales against animal immune systems, as fucose is widely present in the blood.

NagA α-N-acetylgalactosaminidase, NagZ β-N-acetylhexosaminidase (GH109) involved in peptidoglycan recycling occur in multiple copies not only in *Paenibacillus* genome but also in sequenced fungal genomes. It is worth noting that *Peanibacillus* has three times more copies (30) of Nag genes than studied fungi. No proteins from this family had been characterized in Eukaryota.

*Thamnidium elegans* possesses the narrowest set of carbohydrate-processing enzymes which, like in the case of proteases, might be expanded by its associated bacterium. *Paenibacullus* encodes several unique enzymes, like pectate lyases (PL1, PL10, PL9) and rhamnogalacturonan (PL11), oligogalacturonate (PL22) and L-rhamnose-α-1,4-D-glucuronate lyases (PL27). It also encodes lytic chitin monooxygenase (AA10) which may be either used to manipulate the host or to acquire nutrients. Additionally, *Paenibacillus* has expanded digestive repertoire including xylanases and xylosidases (GH11, GH30, GH43) and galactosidases (GH4, GH42, GH110). Interestingly, it encodes bacterial lipid-A-disaccharide synthase (GT19) and murein polymerase (GT51). *Paenibacillus* genome harbours 39 copies of S-layer homology (SLH) domain proteins which are important for biofilm formation [56].

#### Transporters

Sequenced Mucorales have several classes of transporters: Amino Acid-Polyamine-Organocation (APC), Glycoside-Pentoside-Hexuronide (GPH), Putative Tripartite Zn2+ Transporter (TZT), ATP-binding Cassette (ABC), Major Facilitator Superfamily (MFS) and Drug/Metabolite Transporter (DMT) (**Supplementary Table S5)**. *Mucor circinatus* possesses a huge expansion of MFS (347 compared to a median of 209 copies per genome). *Thamnidium elegans* has fewest transporter-encoding genes, but it is again possible that this is an effect of relationship with bacteria serving as a reservoir of necessary enzymes. Both *Umbelopsis* representatives have fewer transporters than analysed Mucorales.

#### Metabolic clusters, secondary metabolites and cofactors

The genome of *Paenibacillus* harbours NRPS, terpene, bacteriocin, T3PKS and S-layer-glycan producing clusters. Mucoromycotina were long considered devoid of secondary metabolite clusters. A review by Kerstin Voigt and colleagues [16] showed genetic determinants for natural product synthesis present in all analysed genomes. Our newly sequenced genomes encode between 3 and 8 secondary metabolite clusters (according to AntiSMASH scans) belonging to different classes. Interestingly, all of the genomes encode terpene clusters (**Supplementary Table S6)** which potentially could produce new natural products like those isolated from *Mortierella* [57].

Umbelopsidales additionally have a trans-2,3-dihydro-3-hydroxyanthranilate isomerase (PhzF) involved in phenazine biosynthesis with yet unknown biological product in fungi [58]. This gene is also present in other fungi and in Endogonales but seems to be missing in Mucorales. Umbelopsidales produce also a citronellol/citronellal dehydrogenase which converts citronellol to citronellic acid an odorous compound with antimicrobial properties. All four Mucorales isolates have a single gene coding for salicylate hydroxylase which is involved in plant host manipulation by *Epichloë* [59] and other fungi.

All analysed genomes have a B12 dependent methionine synthase homologous to the animal variant of the enzyme (PF02965, Met_synt_B12) and B12 independent synthase (PF01717, Meth_synt_2) encountered in Dikarya. Both enzymes have recently been found in other Mucoromycotina [60] and their presence suggests at least partial B12 dependence. Moreover, *Umbelopsis* spp. have methylmalonyl-CoA mutase (MUT), methylmalonyl-CoA/ethylmalonyl-CoA epimerase (MCEE), cob(I)alamin adenosyltransferase (MMAB, pduO) and ribonucleoside-triphosphate reductase (rtpR) - all using cobalamin. Mucorales have a B12-independent ribonucleotide reductase class II (nrdJ) and lack methylmalonyl-CoA mutase and MUT-epimerase genes.Sex locus Sequenced isolates belong to heterothallic genera [64] and genomic screening shows a single sex locus per genome. Generally, these loci contain a single high mobility group (HMG)-domain transcription factor gene (*sexP* or *sexM*), flanked by genes for an RNA helicase (*rnhA*), a triosephosphate transporter (*tptA*) and an alginate lyase (*agl*) [65].

The sex locus of *T. elegans* and *M. plumbeus* is organized like in other Mucorales with genes in the following order *agl/tptA/sex/rnhA* [65]. *M. saturninus* lacks the *tptA* gene and has a sex locus architecture (*agl/sexP/rnhA*) like *M. mucedo* [66]. *M. circinatus* has remains of an integrase instead of the *sex* gene between *tptA* and *rnhA* genes (*agl/tptA/rve/rnhA*). Mobile element insertions in the sex locus have already been documented in *Phycomyces blakesleeanus* [67].

The *rnhA* gene seems to be excluded from the sex locus in both Umbelopsidales genomes, in which the sex locus is organized like in Mucorales but followed by a gene with an additional protein of unknown function belonging to DUF2405 (PF09435) family (*agl/tptA/sexP/DUF2405*). Schultz and colleagues described a similar architecture with the DUF2405 for *Umbelopsis ramaniana* from JGI database [68].

#### Carbon assimilation profiles

Carbon assimilation profiles obtained for six Mucoromycotina strains by screening on Biolog FF microplates are summarized in Supplementary Table (**Supplementary Table S7)**. None of the analysed strains was able to use a full set of 95 tested carbon sources. Each strain was able to grow on 40 to 70 different substrates and had a unique carbon assimilation profile (**Fig.4**) with *T. elegans* being the most versatile degrader (**Fig.5**).

**Figure 4.**
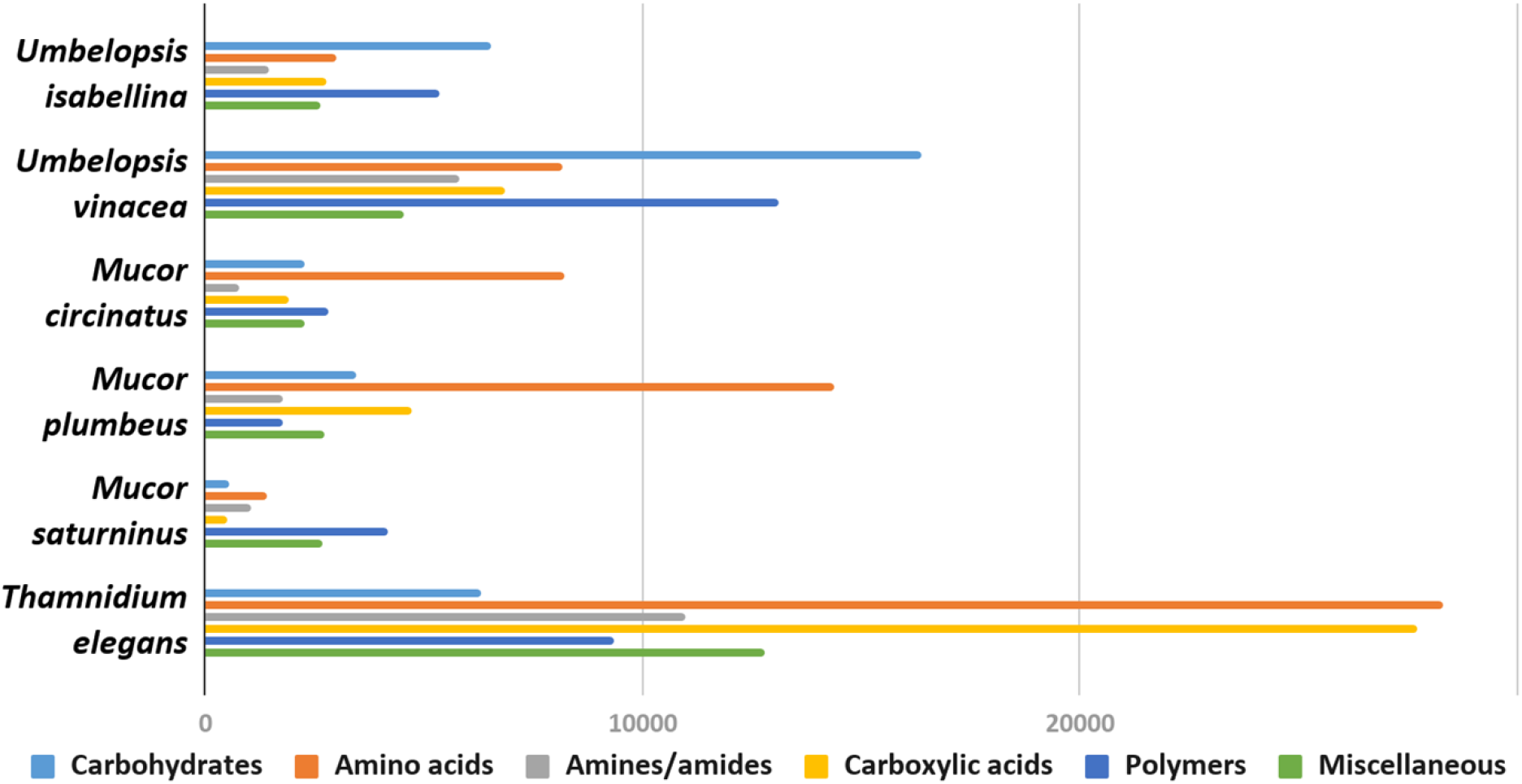
Usage of different types of carbon sources by particular isolates, mean area under the curve (AUC) values aggregated for guilds of carbon sources (according to Preston-et al. [69]).

**Figure 5.**
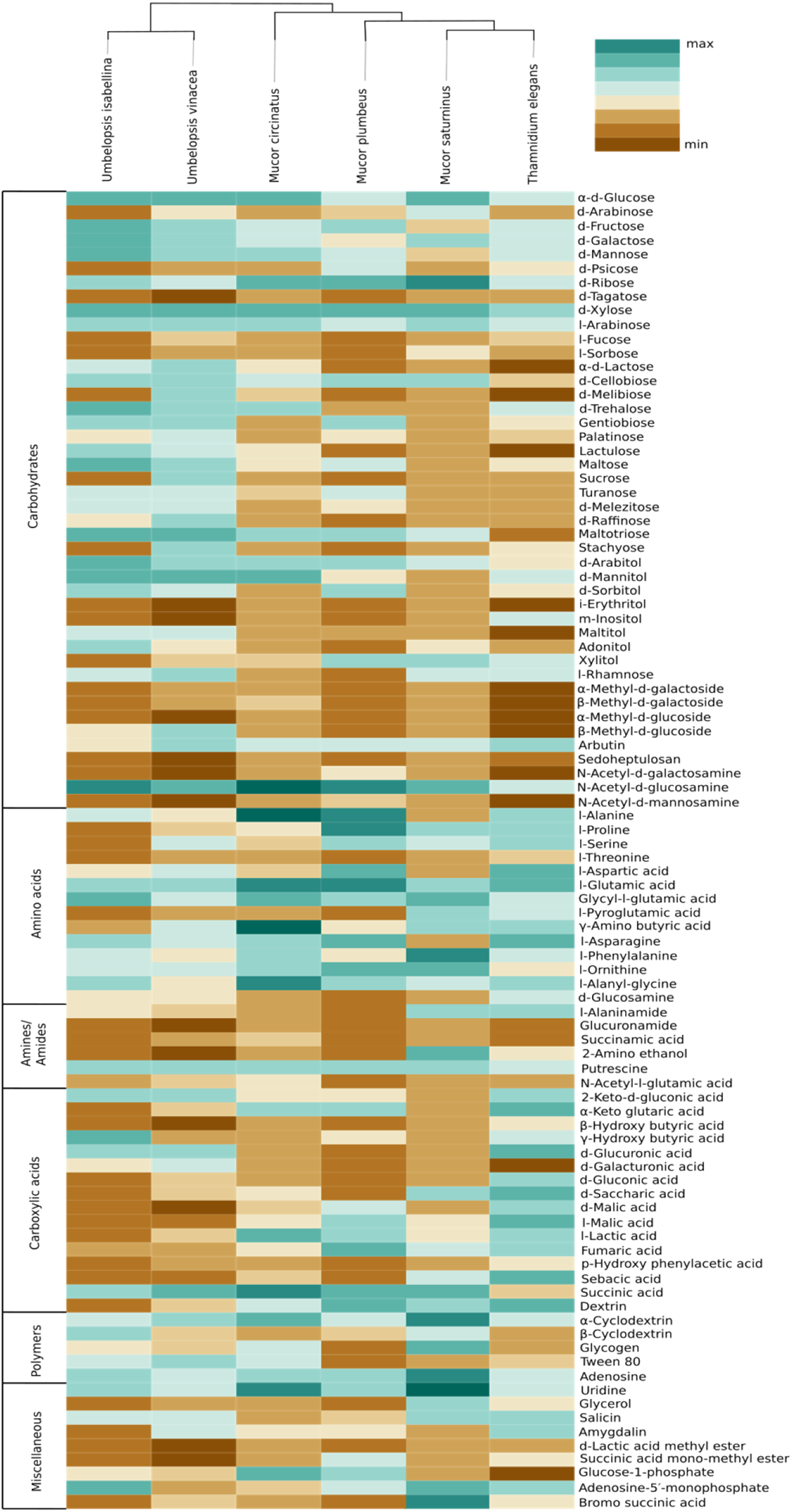
Heatmap representing carbon source utilisation capacity of 6 Mucoromycotina representatives obtained from Biolog FF MicroPlates (scale from brown - not used, to dark green - used very efficiently). The phylogenetic tree is illustrating evolutionary relationships of tested fungal strains. Tested carbon sources are grouped into guilds according to Preston-Mafham et al. [69].

*Umbelopsis* spp. were more efficient in the utilization of carbohydrates, while Mucorales representatives showed the fastest growth rate on amino acids. *Umbelopsis* fungi were able to utilize Adonitol, d-Galacturonic Acid, Maltitol, β-Methyl-D-Glucoside and d-Raffinose whereas none of the four Mucorales grew on these substrates. Additionally, similarly to *Mortierella elongata* AG77 (Uehling et al. 2017), they represented elevated growth rate on other carbohydrates, like d-Galactose, d-Mannose, l-Arabinose, l-Rhamnose, d-Trehalose and lipid (Tween 80), which is consistent with predicted gene models (see the chapter on CaZymes - the presence of rhamnosidase GH78, α-mannosidase GH92, exo-α-L-1,5-arabinanase GH93, α-L-fucosidase GH95). All four *Mucor* species can utilize D-Malic Acid and L-Malic Acid while these substrates seem inaccessible for *Umbelopsis* spp., which have only 3 copies of lactate dehydrogenase whereas *Mucor* spp. have from four to six enzymes from this family. Only *T. elegans* was able to utilize m-Inositol, Sedoheptulosan, β-Hydroxy-butyric Acid, p-Hydroxyphenyl-acetic Acid and L-Threonine. Additionally, *Thamnidium* is distinguished by very efficient development on carboxylic acids. This ability may be explained by the presence of endobacteria whose proteome harbours representatives of M14 and M20 carboxypeptidase families. M14A is present uniquely in both *Umbelopsis* genomes while M14C occurs only in *Paenibacillus*. Also, peptidase T (M20B) is present exclusively in *Paenibacillus*, whereas fungal genomes have several copies of the remaining M20 subfamilies. Some of the analyzed compounds are available only to *Paenibacillus*, based on genomic evidence, e.g. 3-oxoacid CoA-transferase missing from sequenced Mucorales is present in Dikarya and bacteria, including *Paenibacillus*.

#### Fungal lipids

For all six Mucorales, we determined the composition of sterols, fatty acids and phospholipids (**Supplementary Tables S8, S9 and S10)**. The presence of ergosterol, an important component of fungal plasma membranes, contributing to their stability, was confirmed in all 6 fungal biomasses.

Analysed strains contained the following fatty acids: saturated (14:0, 16:0, 18:0), monounsaturated (16:1, 18:1), and polyunsaturated (18:2, 18:3) (**Fig.6, Supplementary Table S8**). The lowest levels were observed for myristic (14:0) and palmitoleic (16:1) acids. A high percentage of oleic acid (18:1) is characteristic for the *Umbelopsis* strains and has also been reported by others [70–72]. It is also the dominant fatty acid for all isolates except for *Thamnidium elegans*. A higher share of linolenic acid (18:3) was observed in Mucorales 18.5% compared to an average of 7.5% in Umbelopsidales. However, the efficiency/quantity of the production/synthesis of particular fatty acids depends, inter alia, on the type of substrate used [73]. Among examined strains, *T. elegans* profile of fatty acids showed the lowest unsaturation index with the domination of palmitic acid (16:0, 31.7%) and the highest share of stearic acid (18:0, 12%), which may suggest increased membrane stiffness and its lower fluidity. Paradoxically, *T. elegans* has more desaturases than the remaining five isolates (6 compared to 4). Production of unsaturated fatty acids involves specialized desaturases. The delta-6-fatty acid desaturase involved in the conversion of 18:2 to 18:3 fatty acid and delta-12-fatty acid desaturase processing 18:1 to 18:2 were experimentally characterized in diverse Mucoromycota including *T. elegans, Mortierella alpina* and *Rhizopus oryzae* [74–76]. All isolates had both delta-6- and delta-12-fatty acid desaturases. *T. elegans* has four copies of delta-12-fatty acid desaturases with an N-terminal short accessory domain of the unknown function (PF11960, DUF3474) which often occurs with desaturase domains and a single delta-6 desaturase with a Cytochrome b5-like Heme/Steroid binding domain (PF00173.28 Cyt-b5).

**Figure 6.**
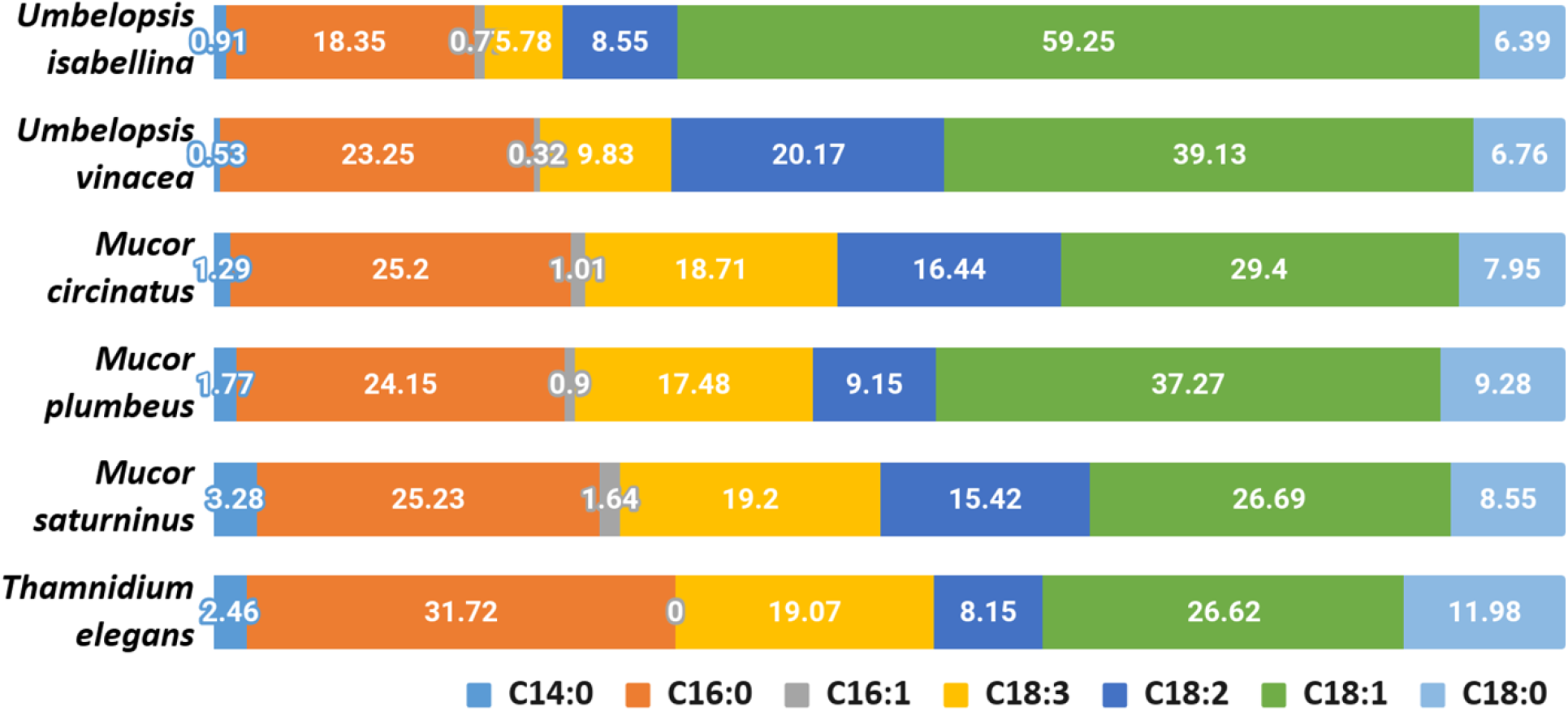
Composition of fatty acids in the analysed strains

Using LC-MS/MS we determined the composition of phospholipids (PLs) present in the biomass of analysed fungi (**Fig.7**, **Supplementary Table S9**). We identified 84 PLs species belonging to 6 classes: phosphatidic acid (PA), phosphatidylcholine (PC), phosphatidylethanolamine (PE), phosphatidylglycerol (PG), phosphatidylinositol (PI), and phosphatidylserine (PS). Sixty-two of them were chosen for subsequent quantitative assessments. This analysis of the PLs for selected strains revealed that PC and PE were predominant for selected Mucoromyotina and constituted up to 56% and 41% of the total cell PLs, respectively. PA, a lipid signal which usually constitutes a minute portion of PLs, in *T. elegans* and *M. saturninus* was found at 8% level. According to several reports, the increased levels of PA in living cells are a consequence of biotic and abiotic stress [77,78]. The open question remains whether *T. elegans* and *M. saturninus* experienced stressing conditions during colony growth in the lab, or the observed increased level of PA is native for them as an adaptation to grow on animal substrate.

**Figure 7.**
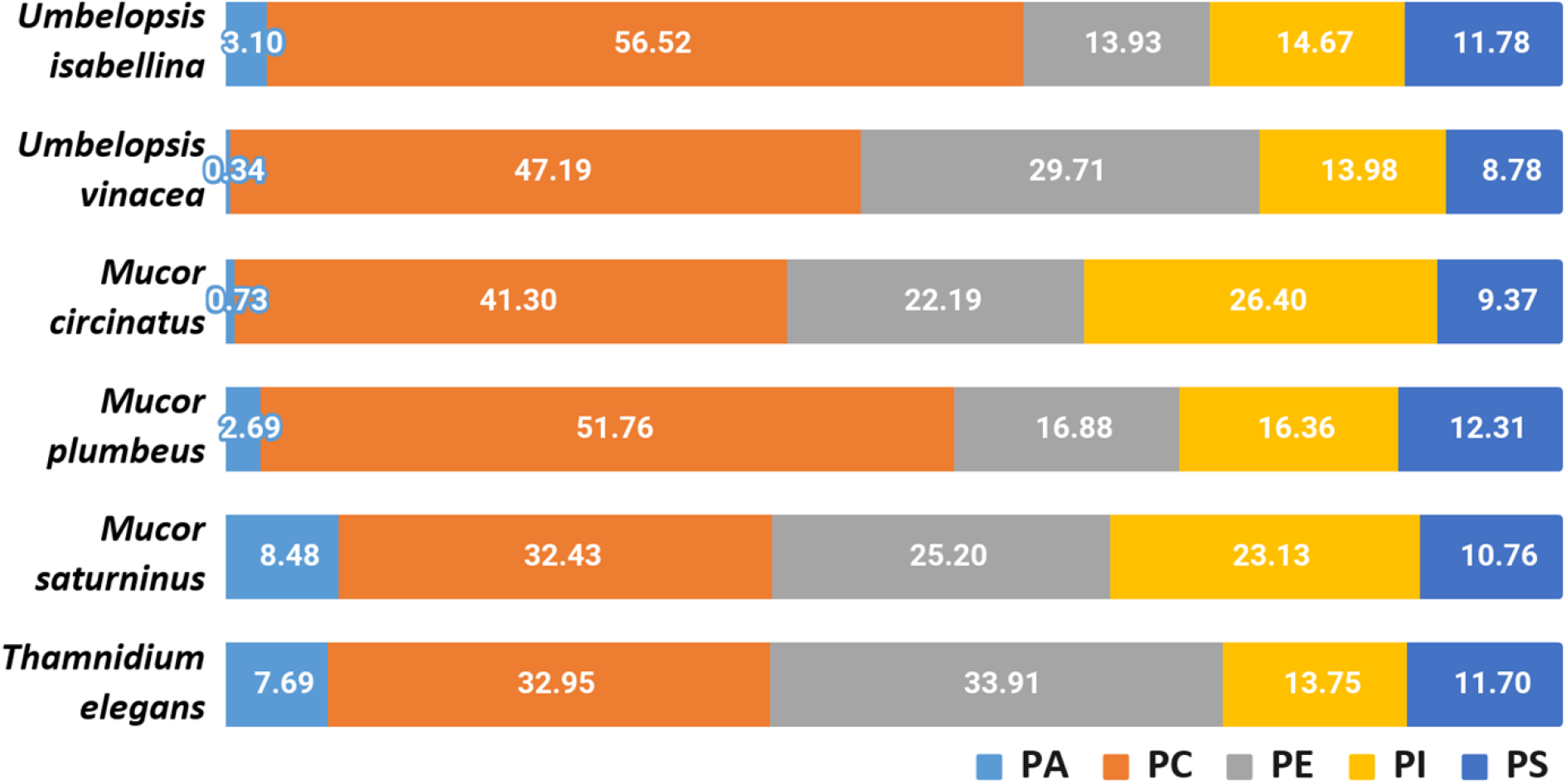
Composition of phospholipids in the analysed strains

An important parameter describing the physical characteristics of biological membranes is PC/PE ratio. PC has a bigger head group than PE. Tighter packing of PE and its acyl chains negatively influences fluidity of membranes as opposed to PC [79]. PC can be synthesized from PE, so they are closely related, and the balance between PE and PC is crucial for maintaining the physiological structure of the cell membrane. Among selected species, *T. elegans* showed the lowest PC/PE ratio (0.79), which may indicate a relatively greater stiffening of the membrane compared to other strains, which is consistent with lowered fatty acids unsaturation index.

Moreover, it is worth mentioning, that only *T. elegans* and *U. vinacea* had traces of PE 16:0 16:1 which is characteristic for diverse bacteria, including isolates of *Pseudomonas putida, Bacillus subtilis* or *Escherichia coli* [80–82]. The dominant PLs species for all selected fungal strains were PC 18:2 18:3, PC 18:1 18:3, PC 18:1 18:1, PE 18:1 18:3, and PE 18:1 18:1. Similar results have already been reported for the PL profile of another Mucoromycotina representative, *Cunninghamella elegans* [78].

Other lipids which are important for the fungal metabolism are acylglycerols, triacylglycerols (TAGs) and diacylglycerols (DAGs) (**Fig.8**, **Supplementary Table S10**). Yeasts store lipids mainly in the form of TAGs, and to less extent, DAGs [33]. In all six fungal strains, both acylglycerols were found. All analysed strains accumulated more TAGs than DAGs but in *T. elegans* this ratio was significantly shifted towards 3:1 TAG to DAG, in contrast to the 5:1 in *M. plumbeus* or even 99:1 in *M. saturninus*. Since TAG synthesis in fungi requires PA and DAG, both present in *T. elegans*, observed DAG/TAG ratio might be a consequence of inhibition of acyl-CoA-dependent diacylglycerol acyl-transferase (DGA1) [83].

**Figure 8.**
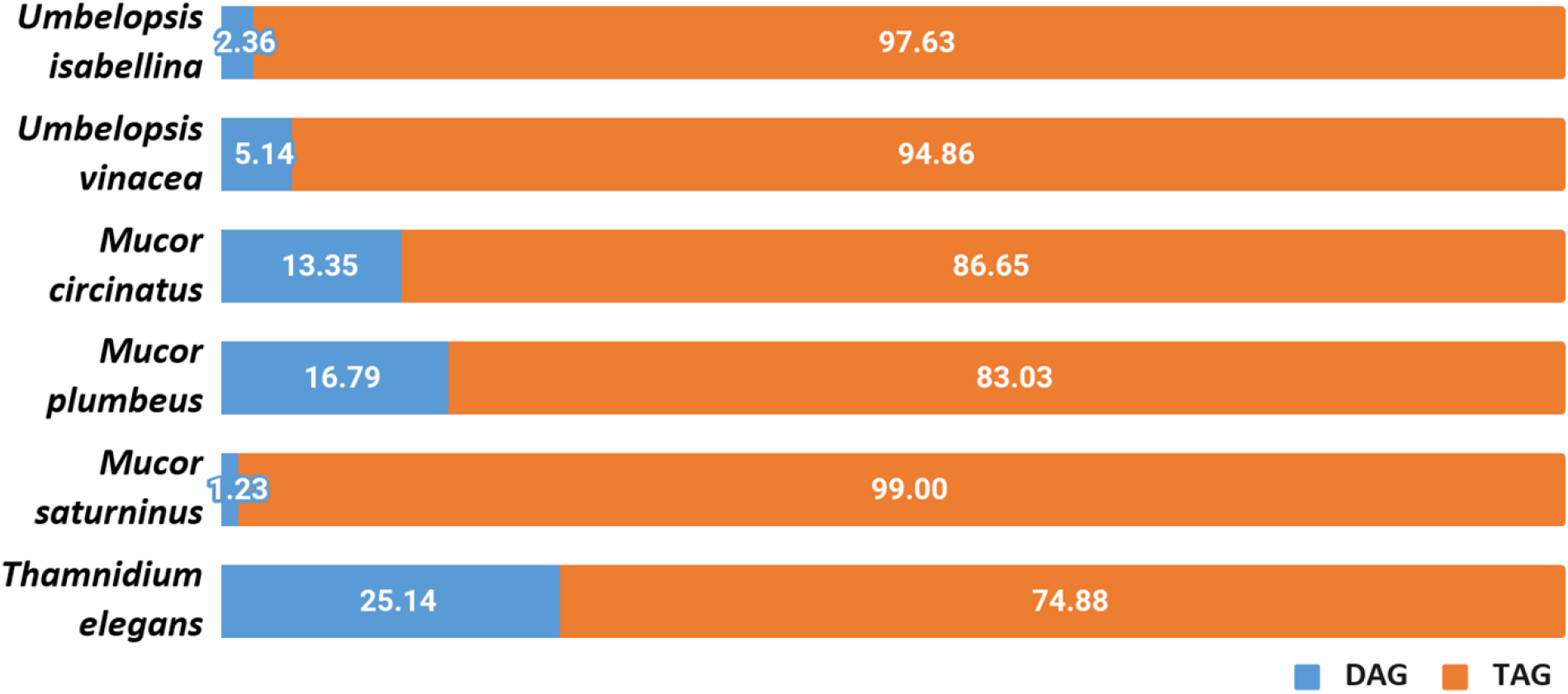
Distribution of diacylglycerols and triacylglycerols in the analysed strains

Lastovetsky and co-workers [33] reported that PE/TAG ratio plays a crucial role in establishing and maintaining symbiosis of the fungus *Rhizopus microsporus* (Mucoromycotina) and its *Mycetohabitans* endobacteria. PE/TAG ratio close to 1:1 is characteristic for symbiosis and departure from that balance might shift the interaction towards antagonism. Interestingly, it was observed that among six strains, the PE/TAG ratio for *T. elegans* was closer to 1:1 (2.21) compared to the average (6.4) for all tested fungi. Analysed strains all contain several copies of diacylglycerol kinase DGK genes (3-5) which is deemed responsible for maintaining the balance between TAG and PE.

The genome of *Paenibacillus* sp. encodes processive diacylglycerol beta-glucosyltransferase required for the synthesis of beta-diglucosyl-DAG - a predominant glycolipid found in Bacillales [84], as well as the Ugp sn-glycerol-3-phosphate transport system which transports glycerol-3-phosphate, essential for phospholipid biosynthesis.

#### Cell-wall carbohydrates

A quantitative analysis of the cell wall carbohydrates revealed the presence of high amounts of glucosamine and fucose, and low amounts of mannose, galactose, and glucose compared to an ascomycetous fungus *Trichoderma reesei* TU-6 (**Fig.9**). Glucosamine content in their cell wall was up to 10 fold higher compared to *Trichoderma*. High glucosamine fraction can be a highlight of the chitin-chitosan cell wall typical for Mucoromycotina [5].

**Figure 9.**
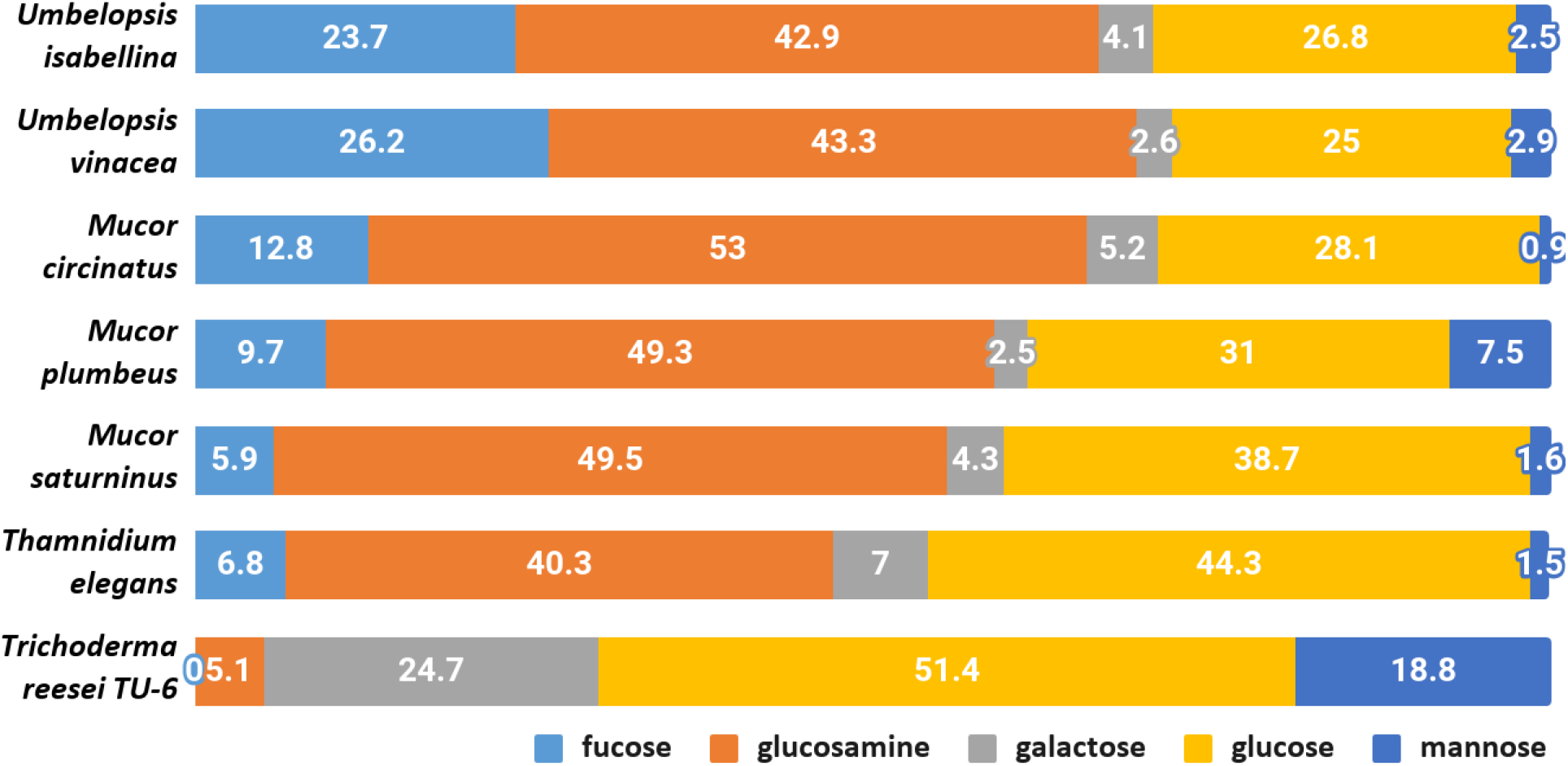
Composition of cell-wall carbohydrates.

Chitin together with glucans participates in the rigidity of the cell wall. The total content of glucosamine (representing chitin/chitosan fraction) and glucose (representing glucan fraction) in four Mucorales: *M. plumbeus, M. saturninus, M. circinatus* and *T. elegans* was above 80% of the cell wall carbohydrates. The amount of these two carbohydrates in *Umbelopsis* strains was lower and reached about 66 and 71% in *U. isabelina* and *U. vinacea*, respectively. *Trichoderma* had only 55% of these sugars in the cell wall and, in addition, glucose was the dominant one.

Melida and coworkers [4] reported that genes encoding chitin biosynthesis (CHS) and its modification (CDA) were present in more copies in *Phycomyces blakesleeanus* and *Rhizopus oryzae*, compared to *Neurospora crassa* (Ascomycota), as well as fewer genes encoding glucan synthases. These results were confirmed in other studies on Mucorales [85] and our results show that respective CaZymes are abundant both in sequenced Umbelopsidales and Mucorales representatives. This could explain the possibility of synthesizing a huge amount of chitin in Mucoromycotina strains compared to *Trichoderma*.

Furthermore, all Mucoromycotina contain fucose, a carbohydrate characteristic for this subphylum, especially in such a high amount. Our study revealed that two *Umbelopsis* strains had from twice to three-fold more fucose when compared to *M. circinatus* and *M. saturninus*, respectively. Presence of fucan in the cell wall was previously reported for *P. blakesleeanus* and *R. oryzae* [4]. During the synthesis of fucan, fucose is transferred from GDP-fucose to polysaccharides by α-fucosyltransferase and this enzyme was detected in the membrane fraction of *M. circinelloides* and partially characterized [85].

The α-fucosyltransferase encoding genes were found in 2 and 4 copies in the genomes of *P. blakesleeanus* and *R. oryzae* respectively, and they were not detected in *N. crassa*, which is in accordance with the fact that no fucose was detected in the cell wall of *Neurospora* [4] and *Trichoderma* (this study). Differences in the copy number of α-fucosyltransferase encoding genes do not explain the significant differences in the amount of fucose in the cell wall of *Mucor* spp. (4-6 copies) and *Umbelopsis* spp. (3-4 copies). All six genomes have 1-3 copies of GDP-L-fucose synthase.

*Thamnidium elegans* despite the associated bacteria shows a cell wall carbohydrate profile similar to remaining Mucorales and other Mucoromycotina. This suggests that the interaction with a bacterial partner alters the metabolism, cell membrane composition but not the exoskeleton of the fungus.

## Discussion

Mucoromycotina display a great diversity of genome sizes and genome complexity [86,87][88–90]. Previously sequenced *Umbelopsis* genomes are very compact with 21-23 Mb and harbour around 9,100 genes (9,081 *U. isabellina* NBRC 7884 [91], *U. isabellina* AD026: 9,193 genes [92] and *U. ramanniana* AG 9,931 genes [92]). Known Endogonales genomes vary between 96 Mb for *Endogone* sp. and 240 Mb for *Jimgerdemannia flammicorona* and harbour from ca. 9,500 to 16,800 genes [25]*. Mucor* and *Rhizopus* species, depending on the number of whole-genome duplications (WGD), have genomes ranging from 25 Mb up to 55 Mb and up to 17,000 genes [54,92,93].

In the present study, we aimed to point several differences between Umbelopsidales and Mucorales taxa. Besides limited genome size, *Umbelopsis* fungi have been demonstrated to be often associated with bacteria and to produce unsaturated fatty acids [70]. Our results build on top of these observations. Sequenced *Umbelopsis* taxa have compact genomes, almost devoid of repetitive sequences, and contain a moderate number of genes. They have a relatively high number of metabolism-related enzymes, especially glycohydrolases compared to Mucorales, but fewer peptidases. Moreover, we showed that all analysed Mucoromycotina display comparable enzymatic capabilities tested on BIOLOG FF microplates and predicted from genomic data regardless of their genome size. Mucorales and Umbelopsidales representatives are mostly soil-inhabiting saprotrophs and require a broad spectrum of secreted enzymes which is reflected in high count of encoded peptidases, glycohydrolases and transporters. Nevertheless, they differ in the ability to use particular carbon sources, produce specific proteases, and have different ratios of different classes of phospholipids.

On one hand, *Umbelopsis* spp. in general have fewer secreted peptidases like pepsins and sublitisins than Mucorales. On the other, they encode several families of metalloproteases and cysteine proteases with unknown functions which are absent from Mucorales. The compact genomes of Umbelopsidales are richer in glycohydrolases and CaZymes. In consequence, they have a tendency to use carbohydrates efficiently and grow fast on these carbon sources. When compared to Mucorales, they produce higher amounts of 18:1 fatty acid and have more fucose in their cell wall. However, the meaning of these findings remains to be understood. Despite sharing an ecological niche, these two orders differ in genome size, associated bacteria and degrading capabilities but, nonetheless, share clear synapomorphic traits. Our experimental results support the validity of 18:3 lipids as a chemotaxonomic marker of Mucoromycotina and fucose as a specific component of their cell wall [4,85].

Previous genomic studies covered diverse Mucorales representatives [41,86,87] whereas Umbelopsidales genomes were published in the form of brief genomic reports [91] without a description of the genomic content. In this study, we aimed to fill this knowledge gap by bringing together genomic analyses with phenotype and biochemical studies, especially in the context of how these fungi function in their environment. Vast carbohydrate related enzyme repertoire observed in Umbelopsidales can be related to their ecology. According to the GlobalFungi database [94] the representatives of the genus *Umbelopsis* are present in 6155 out of 20009 global amplicon samples (ca 30%). They are detected mainly in Europe, North America, and Australia, most often in soil, root, and shoots probes from forest or grassland biomes. The representatives of this genus are well-known late wood colonisers, that probably feed on the substrates which were decomposed by other organisms able to degrade complex substrates like cellulose [95]. However, *Umbelopsis* representatives are also often isolated from living plant material or forest soil [96] and are considered to be plant growth-promoting organisms and root endophytes [97,98]. They can alter plant metabolism leading to the enhanced production of complex metabolites which are not produced without the endophyte [99]. *U. isabellina* and *U. vinacea* described in this study also encoded metabolic clusters including terpenoid clusters.

The representatives of Umbelopsidales were also shown to represent relatively high resistance to some heavy metals like Zn, Mn, Ni or Pb [100] and xenobiotics, such as herbicides [101]. The resistance may be correlated with the presence of numerous ABC transporters encoded in all analysed genomes. Moreover, these genomes have a single arsenite resistance protein homologous to *Absidia repens* BCR42DRAFT_142507 and consisting of ARS2 (PF04959), DUF4187 (PF13821), RRM_1 (PF00076), SERRATE_Ars2_N (PF12066). Such domain architecture is conserved from chytrids to Entomophtoromycotina and Glomeromycotina. Interestingly, the genomes had neither Dikarya-type nor animal metallothioneins.

Umbelopsidales have also been detected in the deep-sea sediments from Magellan seamounts constituting 3.8% of all OTUs [102]. Although this finding is surprising as *Umbelopsis* representatives are well known terrestrial organisms, their presence may be explained by association with plant material. Interestingly, 0.85% of all amplicon samples in which *Umbelopsis* spp. were detected according to GlobalFungi database (Větrovský et al. 2020), are also originating from marine biome. The understanding of this pattern needs further research.

Some representatives of Umbelopsidales are recently considered as effective single cell oils (SCOs) producers as they are capable of producing high amounts of lipids (75% to 84% in dry cell weight; w/w), including polyunsaturated fatty acids (PUFAs). These substances of high dietary and pharmaceutical importance are also considered as precursors for the synthesis of lipid-based biofuels. Although several Mucoromycota representatives were reported to synthesize PUFAs, *U. isabellina* cultivated on glucose has presented exceptionally high lipid production (comparable to the highest values achieved for genetically engineered SCO-producing bacterial strains) [73]. Analysed strains displayed a typical composition of PUFAs with a predominance of oleic acid (18:1), and higher levels of gamma-linolenic acid (18:3) in Mucorales compared to Umbelopsidales. Surprisingly, *Thamnidium* mycelium showed a relatively high saturation index possibly due to interaction with *Paenibacillus*. Bacteria produce more saturated fatty acids during biofilm formation yet it remains to be elucidated if the bacteria-fungus interaction has a similar effect on fatty acid composition in both partners [103]. *Thamnidium* genome contains a high number of lipid metabolism-related genes and there is no clear explanation for the high level of fatty acid saturation. This phenotype was particularly unexpected since *T. elegans* has been used for biotechnological production of gamma-linoleic acid on diverse substrates especially in low temperature [104–106]. It may be hypothesized that an extended set of lipid-processing enzymes in *Thamnidium* is required in order to balance the bacterial impact on lipid homeostasis and shift it back towards fungal characteristics.

Interestingly, several representatives of Umbelopsidales have recently been shown to be colonized by EHB from Burkholderiaceae. Moreover, the metagenomic analysis of another oleaginous fungus - *Mortierella elongata* and its endosymbiont *Mycoavidus cysteinexigens* showed that bacteria alters the metabolism of the fatty acids of the host. Endosymbiont was shown not only to cause declines in the storage of carbohydrates, organic acids and nitrogenous metabolites but also to be involved in the catabolism of fungal fatty acids and changes volatile compounds profiles of the fungus [32]. Among six sequenced Mucoromycotina, we found numerous bacterial reads which assembled into a complete genome only in *Thamnidium elegans*. The presence of an associated bacteria is reflected in, among else, DAG/TAG lipids composition and utilization of carbohydrates which are accessible exclusively to Dikarya and bacteria. This phenomenon together with a limited set of carbohydrate-processing enzymes present in *Thamnidium* excludes the possibility of bacterial contamination and supports the hypothesis of intimate interaction with detected *Paenibacillus*.

Most of the initial reports on intracellular fungal bacteria were based on microscopic observation of uncultivable bacteria inside fungal hyphae [107]. Nowadays, endohyphal bacteria (EHB) can be efficiently identified using genome sequencing methods. The traces of bacterial presence have been found in other published genomes of basal fungi [108]. Identification of EHB can expand our scarce knowledge on the frequency and host range of fungal-bacteria interactions. Our finding of *Paenibacillus* sp. associated with *Thamniudium elegans* is in line with this trend. Diverse *Bacillus* bacteria were found living with truffles [109,110]. *Paenibacillus* has been reported from *Laccaria bicolor* [111], *Sebacina vermifera [112]*, and is known to produce pre-symbiotic and symbiotic interactions with *Glomus [113]*. The genus *Paenibacillus* was erected from *Bacillus* by Ash et al. [114]; it belongs to the family Paenibacillaceae and comprises 253 highly variable species (https://lpsn.dsmz.de/genus/paenibacillus). The representatives of this genus were isolated from a wide range of sources of plant and animal origin. *Paenibacillus* tend to occupy a similar niche to Mucoromycotina moulds as they inhabit soil and dung. The best-studied species - *P. larvae* is known to cause lethal disease of honeybees. However, some other species are known for their plant growth promotion capacities (via siderophores or phytohormones synthesis), others produce a variety of antimicrobials and insecticides. Bacteria from this genus were also shown to produce a plethora of enzymes, like amylases, cellulases, hemicellulases, lipases, pectinases, oxygenases or dehydrogenases [115]. The proteome of *P. larvae* includes a wide range of virulence factors including proteases and toxins [116]. *Paenibacillus validus* stimulated the growth of *Glomus intraradices* [117] and *P. vortex* facilitates dispersal of *Aspergillus fumigatus* [118]. Although the representatives of *Paenibacillus* are known to promote plant growth and secrete several antimicrobial compounds, the endohyphal strain from our study did not encode any antibiotic compounds except for rifamycin-inactivating phosphotransferase. Rather, it provided multiple enzymes that significantly expanded the digestive capabilities of *Thamnidium* while reducing its genome size. Observed genome shrinking cannot be explained solely by the fragmented assembly of *Thamnidium* (its incompleteness is estimated at approximately 5%) because the differences in CaZyme and peptidase abundance from remaining isolates are far greater.

One of the major differences between *Thamnidium* and the remaining isolates is in the lipid composition potentially contributing to cell membrane stiffness. EHB are known to influence host lipid production and bias the TAG/PE ratio [33]. *Thamnidium* had the lowest TAG/PE ratio among tested isolates which might be a sign of symbiotic interaction with *Paenibacillus*, yet the noted ratio was still far from the 1:1 “symbiotic equilibrium” described in Lastovetsky’s report [33]. It is not known whether the TAG/PE values estimated for symbiosis between *Rhizopus microsporus* and *Mycetohabitans* are valid also for other models. It is an open question of how intimate and stable is the interaction between *T. elegans* and *Paenibacillus* sp. The altered traits of *Thamnidium elegans* compared to remaining isolates regardless of their evolutionary distances could be explained by the presence of an associated bacterium classified to *Paenibacillus*. In contrast to lipids, the cell wall carbohydrate composition of *T.elegans* remained unchanged. What we observed is that *Thamnidium* has several metabolic parameters altered but its morphology remained unchanged compared to strains without detectable bacterial partners.

The ancestors of extant Mucoromycotina were present among the first land colonizers and had the ability to access decomposing material. Genome sequencing and phenotyping of Mucorales and Umbelopsidales enabled us to look at the differences of these two old lineages within Mucoromycotina. There are several differences, particularly in their carbon source preferences and encoded carbohydrate repertoire, which hints at subtle niche differentiation. Importantly, predicted digestive capabilities are in line with experimental validation. Early diverging Mucoromycotina possess features characteristic of fungi including ergosterol present in the membranes and a cell wall made of chitin and beta-glucan. Additionally, all studied Mucoromycotina representatives produce 18:3 gamma-linoleic acid and encrust their cell wall with fucose, both of which traits can be a handy discriminant for marking their presence in environmental samples.

## Methods

### Isolates

Six non-pathogenic, common and soil-borne representatives of Mucorales and Umbelopsidales were chosen for sequencing. Two of them represented the *Umbelopsis* genus, isolated from forest soil in Warsaw (Poland). Other two taxa (i.e. *M. circinatus* and *M. plumbeus*) were also soil-derived isolates from both Americas but representing Mucorales order. Finally, two Mucorales representatives that are well known for their proteolytic activity (i.e. *Mucor saturninus* and *Thamnidium elegans*) were also selected (Table 2). Related organisms were tested for biotechnological usage and reference genomic information for some of them (e.g. *U. isabellina*) was already available. The species-level identification of all isolates was confirmed by sequencing of ITS rDNA fragments (according to the protocol proposed by Walther and co-workers [119]) prior to genome sequencing.

**Table 2.** Used strains

| Species | Strain number | Country of isolation | Substrate |
| --- | --- | --- | --- |
| *Umbelopsis isabellina* | WA0000067209 | Poland | forest soil |
| *Umbelopsis vinacea* | WA0000051536 | Poland (Bielany forest) | forest soil |
| *Mucor circinatus* | CBS 142.35 | Brazil | soil |
| *Mucor plumbeus* | CBS 226.32 | Canada | forest soil |
| *Mucor saturninus* | WA0000017839 | Poland (Warsaw) | cheese |
| *Thamnidium elegans* | WA0000018081 | Poland (Truskaw) | horse dung |

### Phenotypic microarray plates

FF phenotypic microarray plates (Biolog Inc., USA) were used to test the capacity of 6 strains to grow on 95 different carbon sources. Carbon sources were grouped into guilds according to Preston-Mafham et al. [69]. All fungal strains were cultured on Potato Dextrose Agar for 7 days and further their swabbed spores were suspended in FF inoculation fluid (deficient amount of carbon sources) to produce a final optical density of 0.036 A at 590 nm. Spores’ suspensions were then inoculated on FF microplates and incubated in the aerobic SpectrostarNano universal plate reader (BMR Labtech, Germany) for 96 h at 20 °C. The analysis of each strain was done in three replicates. The metabolic activity was measured kinetically by determining the colorimetric reduction of a tetrazolium dye. Colorimetric values for wells containing carbon substrates were blanked against the control well. The result was considered positive when a difference between the metabolic activities of the first and last day of incubation was observed in all three repetitions. The mean values and standard deviations of AUC (area under the curve) were calculated for each strain and each guild of carbon sources. The metabolic activity of each species on a particular substrate was represented as a heatmap of log-transformed mean AUC values. All analyses were performed in RStudio using packages ggplot2 and vegan v2.4.2).

### Lipids analysis

#### Extraction

Lipids from fungal cultures of the stationary phase of growth were extracted according to the method proposed by Folch et al. [120] with some modifications. The fungal biomass was filtered and 0.1 mg was transferred into Eppendorf tubes containing glass beads, 0.66 mL of chloroform and 0.33 mL of methanol. The homogenization process using a ball mill (FastPrep) was carried out for 1 min. The mixture was extracted for 2 min. In order to facilitate the separation of two layers, 0.2 mL of 0.9 % saline was added. The lower layer was collected and evaporated. *Phospholipid determination*. The polar lipids were measured using an Agilent 1200 HPLC system (Santa Clara, CA, USA) and a 4500 Q-TRAP mass spectrometer (Sciex, Framingham, MA, USA) with an ESI source. For the reversed-phase chromatographic analysis, 10 μL of the lipid extract was injected onto a Kinetex C18 column (50 mm × 2.1 mm, particle size: 5 μm; Phenomenex, Torrance, CA, USA). The mobile phase consisted of 5-mM ammonium formate in water (A) and 5-mM ammonium formate in methanol (B). The solvent gradient was initiated at 70% B, increased to 95% B over 1.25 min, and maintained at 95% B for 6 min before returning to the initial solvent composition over 3 min. The column temperature was maintained at 40 °C, and the flow rate was 500 μL min-1. The instrumental settings of mass spectrometer were as follows: spray voltage −4500 V, curtain gas (CUR) 25, nebulizer gas (GS1) 50, turbo gas (GS2) 60, and ion source temperature of 600 °C. The data analysis was performed with the Analyst™ v1.6.2 software (Sciex, Framingham, MA, USA).

Two approaches were applied to identify PLs: targeted and untargeted. The untargeted approach was performed with the precursor ion scanning (precursor for m/z 153) survey scan, triggering the EPI experiments. On the basis of the untargeted analysis, a comprehensive list of the multiple reaction monitoring (MRM) transitions was generated.

#### Acylglycerols

DAGs and TAGs analysis was undertaken by liquid chromatography coupled to mass spectrometry (LC-MS) with electrospray ionization (ESI) on an QTRAP 4500 (Sciex). A Kinetex C18 column (see phospholipids determination) and mobile phases consisting of water (A), a mixture of acetonitrile:isopropanol (5:2, v/v) (B) and 5 mM ammonium formate with 0.1 % formic acid were used. The solvent gradient was initiated at 35% B, increased to 100% B over 4 min, and maintained at 100% B for 11 min before returning to the initial solvent composition over 2 min. The column temperature was maintained at 40 °C, and the flow rate was 600 μL min-1. The QTRAP 4500 was operated with positive ionization at an electrospray voltage of 5500 V and a targeted multiple reaction monitoring (MRM) approach containing transitions for known precursor/product mass-to-charge ratio. Under these conditions, the TAG ionize as ammonium adducts.

#### Fatty acid analysis

A lipid sample was diluted in 1.5 mL of methanol and transferred to a screw-capped glass test tube. To the lipid solution, 0.2 mL of toluene and 0.3 mL of the 8.0% HCl solution were added [121]. The tube was vortexed and then incubated at 45 °C overnight. After cooling to room temperature, 1 mL of hexane and 1 mL of water were added for the extraction of fatty acid methyl esters (FAMEs). The tube was vortexed, and then, 0.3 mL of the hexane layer was moved to the chromatographic vial. 1.6 μL of the extract samples were analyzed using gas chromatography.

A FAMEs analysis was performed with an Agilent Model 7890 gas chromatograph, equipped with a 5975C mass detector. The separation was carried out in the capillary column HP 5 MS methyl polysiloxane (30 m × 0.25 mm i.d. × 0.25 mm ft). The column temperature was maintained at 60 °C for 3 min, then increased to 212 °C at the rate of 6 °C min-1, followed by an increase to 245 °C at the rate of 2 °C min-1, and finally to 280 °C at the rate of 20 °C min-1. The column temperature was maintained at 280 °C for 10 min. Helium was used as the carrier gas at the flow rate of 1 ml min-1. The injection port temperature was 250 °C. The split injection was employed. Fungal fatty acids were identified by comparison with the retention times of the authentic standards (Sigma, Supelco) and the results were expressed as a percentage of the total amount of fatty acids.

*Sterol analysis* was undertaken using the QTRAP 3200 (Sciex) mass spectrometer connected to a 1200 series HPLC system. A Kinetex C18 column (see phospholipids determination) was used. The solvents were: water and methanol, both containing 5mM ammonium formate. Analytes were eluted with the following gradient: 40% solvent B from 0 to 1 minute, 100% solvent B from 1 to 4 minutes, 40% solvent B from 4.0 to 4.1 minutes, 40% solvent B from 4.1 to 6 minutes with the flow rate 0.8 ml min^−1^. The QTRAP instrument was set to the positive ion mode, with the atmospheric pressure chemical ionization (APCI) temperature of 550°C.

### Cell wall preparation and determination of cell wall carbohydrates

Fungi were cultivated in PDB medium, washed with 10 mM Tris/HCl, pH 7.5, suspended in the same buffer, disintegrated with 0.5 mm glass beads in the presence of a protease inhibitor cocktail (Sigma-Aldrich) and centrifuged at 1500 x g for 10 min. The resulting pellet containing cell walls was washed with ice-cold 1 M NaCl until the disappearance of absorbance at 260-280 nm [122]. The lyophilized cell wall was hydrolyzed o/n in 4 M trifluoroacetic acid (TFA) at 100° C. After cooling on ice, samples were centrifuged at 17 000 x g for 5 min at 4°C. The supernatant was dried under N2 and washed twice with pure methanol. After removing methanol with N2, the pellet was resuspended in Mili Q water and purified on a Millipore Filter Device (0.45 μm pores) by centrifugation at 16 000 x g for 4 min. Samples were stored at −20°C. Monosaccharides were determined by high-performance anion-exchange chromatography using a Dionex ICS-3000 Ion Chromatography System with a Carbo Pac PA10 analytical column. Neutral sugars were eluted with 18 mM NaOH at 0.25 ml/min [123].

### Culture conditions and DNA extraction

All fungal strains were cultured on 4% Potato Dextrose Agar for 7 days at 20°C. Total genomic DNA was extracted from 30 mg of fresh mycelium following a CTAB-based chloroform extraction protocol [124] and cleaned-up following caesium chloride density gradient centrifugation method [125]. DNA quality and concentration were estimated by 1% agarose gel electrophoresis and NanoDrop^®^ (Thermo Fisher Scientific). Quantification of purified DNA was performed on a Qubit Fluorometer. The identity of all strains was confirmed by the preliminary sequencing of the internal transcribed spacer (ITS rDNA) region and with standard morphological identification procedures.

### Sequencing

DNA was sequenced at the High Throughput Sequencing Facility of UNC, Chapel Hill, NC, USA.

Whole-genome sequencing was accomplished using a hybrid approach, combining Illumina short-read data with PacBio long-read data.

Total cellular DNA was sheared using a Covaris E220 sonicator to achieve fragments with an average size of 500 bp. Then libraries were prepared using the Kapa Hyper kit. Libraries were size selected for insert fragments around 500 base pairs using Pippin Prep automatic DNA size selection system (Sage Science). Libraries were analyzed and quantified using a LabChip GX automated electrophoresis system (Caliper) and pooled. The pools were sequenced on Illumina MiSeq sequencer paired-end sequencing (2×300 cycles) to obtain longer reads and on Illumina HiSeq sequencer 2500 (2×150 cycles) to obtain more coverage.

For the PacBio RSII data, ten microgram aliquots of genomic DNA were sheared in a Covaris g-TUBE to a target fragment size of 20 kb using the shearing conditions provided in the Covaris g-TUBE user manual. The protocol for preparing a 20 kb library (Pacific Biosciences Procedure and Checklist-20kb Template Preparation Using Blue Pippin™ Size-Selection system) was subsequently followed, using 5 μg of purified, sheared DNA as starting material. Template concentration was calculated using the Qubit fluorometer and the average size was determined by BioAnalyzer trace analysis and served as input to the Annealing & Binding Calculator v.2.1.0.2 (Pacific Biosciences) to prepare SMRTbell-template annealing and polymerase-template binding reactions, as well as the final dilution of the polymerase-bound template complex for sample plate loading and spike-in of control DNA. The PacBio reads were filtered to a minimum read length of 100 bp and a minimum read quality score of 0.85.

### Assembly

PacBio reads were processed with SMRT Analysis Software. Spades (v.3.10.1) [126] was chosen as a slow but high-quality assembler [127]. An initial assembly of Illumina reads alone with Abyss [128] produced very fragmented and incomplete assemblies. Quast (v.4.6.1) was used to select the best assembly [129]. The completeness of genome assembly and gene annotation was assessed using BUSCO (v.3b) [38], using BUSCO’s fungal dataset universal single-copy orthologs from OrthoDB9 as reference.

BlobTools2 [130] was used to partition the scaffolds, on the basis of read coverage, G/C content, and taxonomic affiliation. The bacterial and fungal reads were assembled separately.

### Gene calling

MAKER (v.2.31.8) [131] was used as the main annotation pipeline for gene prediction and annotation. As part of MAKER, AUGUSTUS (3.3) [132], and GeneMark-ES fungal version (v.4.21) were used as *ab initio* gene predictors. The *ab initio* was trained for a couple of times within MAKER based on sequence similarity search via BLAST and sequence alignment via Exonerate (v.2.2) [133] with default parameter settings, using 100,082 EST and 273,458 protein sequences of Mucoromycotina obtained from NCBI in January 2018, together with SwissProt reference proteins (downloaded 18th January 2018). Repeats were predicted with RepeatMasker (v.4.0.7) [134] together with Tandem Repeat Finder (TRF) (v.4.04) [135] and RMBlast (NCBI blast package of RepeatMasker), based on Repbase database 2017 edition [136] as described in [39].

Gene calling in Paenibacillus was performed with Prokka [137], an antimicrobial component search was done in Abricate [138]. A whole genome-based taxonomic analysis was performed in Type (Strain) Genome Server (TYGS) [139] on 2020-06-01.

### Protein coding gene annotation

Predicted protein-coding genes were mapped on Cazy with DBCan [140] modified like in [141], MEROPS protease evalue 1E-10 [142], PFAM evalue 0.001 [143], INTERPROScan 5.28-67 (v[144]), CDD evalue 1E-5 [145], AntiSMASH 4.0 (--transatpks_da --clusterblast --subclusterblast --knownclusterblast --smcogs --cassis --borderpredict --full-hmmer) [146], TCDB evalue 1E-5 [147]. RNAi components were filtered with an evalue of 1E-10. KEGG mapping was performed with kofamscan 1.3.0 with default settings [148].

### Phylogenetic analysis of bacteria found in the genome of Thamnidium

16S rRNA sequence was extracted from the bacterial reads found in Thamnidium using blast. Then it was combined with publicly available 16S sequences of several species and strains of *Paenibacillus* and *Bacillus* (GB accession numbers can be found on Fig. 1). Sequences were then aligned using MAFFT [149] and trimmed using trimal - automated1 [150]. Then the best evolution model was detected using modeltest-ng across all evolutionary models [151] (TrN+I+G4 model selected based on AIC, BIC and AICc criteria) and phylogenetic tree was calculated using raxml-ng [152] with 1000 bootstrap replicates. The tree was then rooted using four *Bacillus* sequences.

## Acknowledgements

We thank Gustavo Henrique Goldman and Marcin Grynberg for their insight and comments about the manuscript. This work was supported by the National Science Centre, Poland, under grants no 2012/07/D/NZ2/04286 and 2017/25/B/NZ2/01880 to Anna Muszewska, 2019/35/D/NZ2/03411 to Kamil Steczkiewicz and 2015/17/D/NZ8/00778 to Julia Pawlowska.

The funders had no role in study design, data collection and analysis, decision to publish, or preparation of the manuscript.

## Data Availability Statement

All of the genomic data have been deposited in the NCBI BioProject PRJNA668042. Raw reads for all fungal strains are available in the SRA database under accession numbers SRR12875449-SRR12875464. Assemblies and annotations are deposited under accession numbers XXXXXXX-XXXXXX. Bacterial reads, assembly, and annotation can be found under accession numbers SRAXXXX, XXXXX, and XXXX, respectively.

The sequence analyses are based on publicly available sequences from NCBI and UNIPROT.

## Author Contributions

A.M. and J.P. designed the study, A.M. performed assembly, annotation and sequence analyses, J.S.K, P.B., A.O., K.S., J.P. and A.M. interpreted the data and drafted the manuscript, K.S., J.P. and A.M. wrote the manuscript, E.M. and P.M. performed whole-genome sequencing, O.S., U.Z. and S.P. performed experimental procedures and prepared the samples, T.A-P and K.SA. performed carbon source usage experiments U.P-L and J.S.K. analysed cell wall composition, P.B. analysed lipid composition, J.P. analysed carbon source usage.

## Supplementary Materials

**Supplementary Figure S1.** Tree inferred with FastME 2.1.6.1 [153] from Genome BLAST Distance Phylogeny approach distances calculated from genome sequences of Paenibacillus isolates [154]. The numbers above branches are pseudo-bootstrap support values > 60 % from 100 replications, with average branch support of 47.8 %. The tree was rooted at the midpoint by the program. Pae_T is the *Paenibacillus* sequenced as associated with *Thamnidium elegans* groups with *P. illinoisensis* strains.

## Supplementary Tables

**Supplementary Table S1** Repetitive elements in analysed genomes

**Supplementary Table S2** RNAi pathway components in the analysed genomes

**Supplementary Table S3** Merops families of peptidases in analysed genomes

**Supplementary Table S4** CaZyme families in analysed genomes

**Supplementary Table S5** TCDB families of transporters in analysed genomes

**Supplementary Table S6** Antismash metabolic clusters in analysed genomes

**Supplementary Table S7** Carbon source usage of the isolates

**Supplementary Table S8** Total lipid and fatty acid composition of the isolates

**Supplementary Table S9** Phospholipid composition of the isolates

**Supplementary Table S10** Diacyglicerols and triacyglicerols content in the analysed isolates

**Supplementary Table S11** TYGS results for *Paenibacillus* sp. genome listing DDH scores for related organisms.

